# Clinical Knowledge and Reasoning Abilities of AI Large Language Models in Pharmacy: A Comparative Study on the NAPLEX Exam

**DOI:** 10.1101/2023.06.07.544055

**Authors:** Mirana Angel, Anuj Patel, Amal Alachkar, Pierre Baldi

## Abstract

**Objective:** This study aims to evaluate the capabilities and limitations of three large language models (LLMs) – GPT-3, GPT-4, and Bard, in the field of pharmaceutical sciences by assessing their pharmaceutical reasoning abilities on a sample North American Pharmacist Licensure Examination (NAPLEX). We also analyze the potential impacts of LLMs on pharmaceutical education and practice.

**Methods:** A sample NAPLEX exam consisting of 137 multiple-choice questions was obtained from an online source. GPT-3, GPT-4, and Bard were used to answer the questions by inputting them into the LLMs’ user interface. The answers provided by the LLMs were then compared with the answer key.

**Results:** GPT-4 exhibited superior performance compared to GPT-3 and Bard, answering 78.8% of the questions correctly. This score was 11% higher than Bard and 27.7% higher than GPT-3. However, when considering questions that required multiple selections, the performance of each LLM decreased significantly. GPT-4, GPT-3, and Bard only correctly answered 53.6%, 13.9%, and 21.4% of these questions, respectively.

**Conclusion:** Among the three LLMs evaluated, GPT-4 was the only model capable of passing the NAPLEX exam. Nevertheless, given the continuous evolution of LLMs, it is reasonable to anticipate that future models will effortlessly pass the exam. This highlights the significant potential of LLMs to impact the pharmaceutical field. Hence, we must evaluate both the positive and negative implications associated with the integration of LLMs in pharmaceutical education and practice.

## Introduction

In recent years, the field of Artificial Intelligence (AI) has witnessed remarkable advancements, propelled by breakthroughs in machine learning, deep neural networks, and the rapid development of modern computational devices [1]. AI’s increasing accuracy has led to numerous applications in the healthcare industry, including disease diagnosis, virtual assistants, remote monitoring, and precision medicine. [2]. Moreover, AI-based large language models (LLMs), which undergo training using extensive collections of textual data, possess the remarkable ability to effortlessly produce top-notch text (as well as software) and present novel prospects for revolutionizing the field of healthcare. But how knowledgeable and capable of reasoning are LLMs when it comes to pharmaceutical sciences? While previous studies have accessed LLMs’ capabilities in the clinical setting [3], here we access their capabilities in a more chemical and drug-related discipline.

Large language models employ sophisticated deep-learning techniques to process and generate human-like text. These models typically rely on transformer architectures, which utilize the attention mechanism to effectively capture and utilize contextual information [4, 5]. The success of large language models can be attributed to their ability to capture complex patterns and dependencies in text data, enabling them to generate coherent and contextually relevant responses. For our experiment, we consider the most notable LLMs, including GPT-3, GPT-4, and Bard, which are built based on the decoder-only architecture of the transformer. [6, 7, 8]

Specifically, GPT-3 contains over 175 billion parameters and has the capability to complete diverse tasks, whereas GPT-4 is a much bigger model that contains one trillion parameters. [6, 7]. Bard is a Google Chatbot that is initially built based on the 137-billion-parameter LaMDA model [9] but then switched to the PaLM-2 model with 540 billion parameters [8]. With this new update, we expect Bard to have better pharmaceutical reasoning ability than GPT-3, although it still may be shy of GPT-4.

To further assess the capabilities and limitations of the aforementioned LLMs in the healthcare industry, more specifically for pharmacy, we analyze their performance on The North American Pharmacist Licensure Examination (NAPLEX). Passing the NAPLEX is a requirement for obtaining one’s pharmacy license in all 50 states in the United States. The exam is divided into three core sections: managing drug therapy, accurately and safely preparing and dispensing medications, and promoting public health [10, 11]. Successfully passing this examination signifies an individual’s aptitude and expertise in pharmacy practice.

## Materials and Methods

In our study, we employed an assessment comprising a set of 137 multiple-choice questions sourced from a representative test repository designed to evaluate the requisite knowledge and proficiencies essential for the standard NAPLEX examination. Out of the total 137 multiple-choice questions, 28 questions necessitated the selection of multiple answers to obtain complete credit, while 1 question demanded a numerical response and 1 question required a true or false declaration [12].

In our experiment, we inputted all 137 multiple-choice questions into GPT-3 and GPT-4 via the ChatGPT user interface [13], while using the Google-provided user interface for Bard [3]. Each question was entered individually into the interface, and the corresponding response was recorded. After capturing all the answers, we compared them to the answer key provided by the website and assigned grades based on the accuracy of the responses.

## Results

From the assessment results, GPT-3 and Bard scored 51.1% and 67.8% questions correct respectively. GPT-4, although was able to score much higher than both, answering 80.2% of the questions correctly (Table 1).

**Table 1:** The overall accuracy of each LLM on the NAPLEX exam

| Model | Raw Score | Percentage Correct |
| --- | --- | --- |
| Bard | 93/137 | 67.8% |
| GPT-3 | 70/137 | 51.1% |
| GPT-4 | 108/137 | 78.8% |

Upon evaluating the relationship between the parameter size of each Language Model and its corresponding accuracy in answering questions on the NAPLEX exam, our analysis reveals a positive correlation. Notably, larger model sizes tend to exhibit higher scores. Specifically, GPT-4, boasting an impressive 1 trillion parameters, achieved the highest performance, while GPT-3, with a comparatively modest parameter count of 175 billion, attained the lowest score (Figure 1).

**Figure 1:**
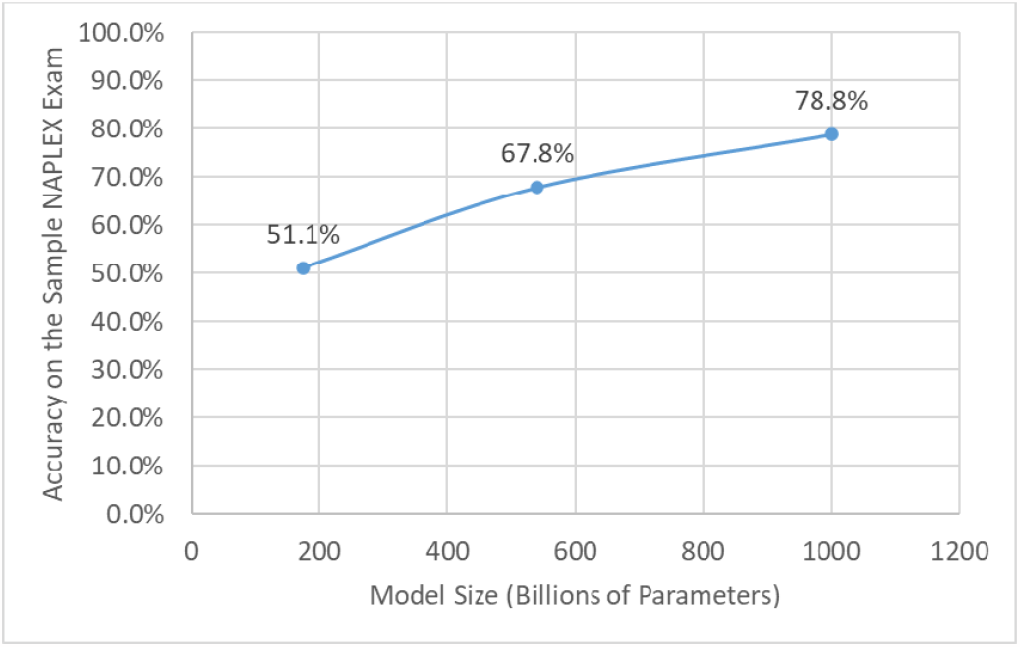
The relationship between the size of the LLM and its performance on the sample NAPLEX exam

When focusing solely on single-answer questions within the exam, each Language Model demonstrated improved performance compared to its overall score. Specifically, Bard achieved a notable score of 82.2%, indicating a substantial increase in accuracy. Similarly, GPT-3 displayed enhanced performance with a score of 62.6%, while GPT-4 exhibited the highest accuracy among the LLMs with an impressive score of 87.9% on single-answer questions (Table 2).

**Table 2:** Test results of different models on the same assessment when comparing only questions involving a single answer.

| Model | Percentage Score |
| --- | --- |
| Bard | 82.2% |
| GPT-3 | 62.6% |
| GPT-4 | 87.2% |

When focusing on multiple-answer questions, the LLM’s tested poorly compared to overall and single-answer questions. Out of the 28 multiple answer questions, Bard answered 21.4% questions correctly, GPT-3 scored 17.9%, and GPT-4 scored 53.6% (Table 3).

**Table 3:** Test results of different models on the same assessment when comparing questions requiring multiple answers.

| Model | Percentage Score |
| --- | --- |
| Bard | 21.4% |
| GPT-3 | 17.9% |
| GPT-4 | 53.6% |

Additionally, the LLMs were unable to understand or answer some of the questions due to limitations such as the lack of knowledge of recent events and restrictions on answering sensitive health-related questions. As a result, Bard could not understand 14 questions, while GPT-3 could not answer 2 questions and GPT-4 could answer all questions (Table 4).

**Table 4:** Number of questions unable to be answered by each LLM due to limitations

| Model | Questions Unable to Answer |
| --- | --- |
| Bard | 14 |
| GPT-3 | 2 |
| GPT-4 | 0 |

## Discussion

The determination of the passing score for the NAPLEX (North American Pharmacist Licensure Examination) is undertaken by the National Association of Boards of Pharmacy (NABP). The NABP employs a criterion-referenced scoring process that considers the varying difficulty levels of different examination forms and aims to ensure fairness to all candidates. The passing score is initially established as a raw score, which is then subjected to mathematical transformation to yield a scaled score. This transformation is performed to ensure that the minimum passing scaled score is set at 75. It is important to note that achieving a 75% accuracy does not necessarily equate to passing the exam, as the difficulty of questions may vary across different examination forms. However, it can generally be considered a reasonable indicator of a passing score on the actual NAPLEX. [14]

The results presented in Table 1 provide insights into the performance of different language models, namely GPT-3, Bard, and GPT-4, in relation to their overall accuracy on the sample NAPLEX. GPT-3 achieves an accuracy of 51.1%, indicating its limitations in answering NAPLEX questions accurately. Bard, on the other hand, demonstrates a higher accuracy of 67.8%, showcasing some improvement in performance compared to GPT-3. However, neither of these models meets the 75% accuracy threshold required to pass the sample exam. In contrast, GPT-4 exhibits a significantly enhanced performance, surpassing the passing threshold with an overall accuracy of 78.8%. These findings suggest that larger language models have higher capabilities in addressing NAPLEX questions, and as model sophistication continues to evolve, it is reasonable to anticipate even higher levels of accuracy, potentially approaching 100%. However, Figure 1 reveals a concave relationship between accuracy and model size, indicating that achieving further improvements in accuracy may necessitate the use of significantly larger models. This implies that while the advancements in model development have demonstrated promising outcomes, future enhancements may require exponential increases in model size to continue improving the accuracy.

In addition to analyzing the performance of LLMs in relation to overall accuracy, a closer examination of specific question types sheds light on their abilities to address different formats within the NAPLEX. Table 2 presents the scores achieved by GPT-3, Bard, and GPT-4 when considering multiple-choice questions with only one correct answer. GPT-3 obtains a score of 62.6%, Bard achieves a higher score of 82.2%, and GPT-4 demonstrates superior performance with a score of 87.2%. However, when focusing on questions with multiple correct choices (select all answers that apply), Table 3 reveals lower accuracy rates. GPT-3 scores 17.9%, Bard achieves 21.4%, and GPT-4 demonstrates the highest accuracy at 53.6%. These findings indicate that while the models perform reasonably well on multiple-choice questions with a single correct answer, their accuracy declines when faced with the challenge of questions involving multiple correct choices. This suggests that further improvements and considerations may be necessary to enhance the model performance in tackling more complex question formats within the NAPLEX. The results presented in Table 4 offer valuable insights into the limitations of Language Models (LLMs) in comprehending pharmaceutical problems. During the evaluation of LLMs using the NAPLEX model examination, Bard exhibited difficulty in answering a subset of questions, providing a response such as “*I’m not programmed to assist with that*.” In a similar vein, GPT-3 faced challenges in retrieving pertinent information for more recent queries and responded with the following: “*Unfortunately, as of my knowledge cutoff in September 2021, there is no information available about a drug called Conjupri being recently approved by the FDA. Therefore, I cannot provide information regarding its pharmacologically active enantiomer*.” Conversely, GPT-4 demonstrated proficiency in responding to all the problems posed. Despite both GPT-3 and GPT-4 sharing the same cutoff date of September 2021 [20], GPT-4 showcased an ability to locate relevant data related to questions concerning recent information. This enhanced performance can likely be attributed to its substantially larger parameter size, enabling it to access less popularized but pertinent data at the time. In contrast, Bard’s inability to comprehend 14 questions provides valuable insight into the capabilities and limitations of Bard’s updated model employing a PaLM-2 framework.

The inability of these LLMs to locate recent data or comprehend pharmaceutical-related questions poses a problem. In the context of time-sensitive matters like emerging diseases such as COVID-19, access to up-to-date and relevant information is crucial for harnessing the full potential of LLMs. If an LLM fails to understand a pharmaceutical-related question that relies on pertinent and recent information, its utility to scientists in the field becomes limited. Addressing this issue is essential to ensure that LLMs can effectively assist researchers and scientists in the ever-evolving landscape of the pharmaceutical domain.

The advent of large language models (LLMs) has had significant implications for pharmacy education. On the one hand, LLMs offer numerous benefits by granting students access to vast amounts of information and resources, which enables them to enhance their understanding of pharmaceutical concepts and improve their problem-solving skills. These models can serve as powerful tools for educators, helping them create engaging learning experiences. Moreover, LLMs can support personalized learning, allowing students to receive tailored feedback and guidance. [14, 15] However, educators must be aware of potential negative impacts. Overreliance on LLMs may lead to a passive learning approach, potentially diminishing critical thinking and analytical abilities. [16] To mitigate these negative effects, educators should emphasize the importance of critical thinking and promote active student engagement. They should also guide students in assessing the credibility and reliability of information obtained from LLMs, thus fostering information literacy skills. Furthermore, educators should adopt a more personal and interactive approach to evaluating student learning outcomes, rather than relying solely on multiple-choice exams and text-based assessments. In cases where a multiple-choice exam is necessary, educators can opt for a format that includes multiple correct answers, as this approach better evaluates student learning.

The impact of LLMs extends beyond pharmacy education and has far-reaching implications for the practice of pharmacy and pharmaceutical sciences. LLMs have the potential to revolutionize medication management and patient care by providing real-time access to comprehensive drug information, potential drug interactions, and personalized treatment plans. This can lead to improved patient outcomes and enhanced medication safety. Moreover, LLMs can significantly promote pharmaceutical research, enabling more efficient data analysis, knowledge discovery, and hypothesis generation. [17] By leveraging LLMs, researchers can access vast amounts of scientific literature, accelerating the process of literature review and facilitating the identification of new research directions. However, the integration of LLMs may also lead to workforce changes and job displacement, as certain tasks traditionally performed by human professionals may become automated. [18] It is crucial for the pharmacy profession to adapt to these changes by embracing interdisciplinary collaborations, developing skills in data interpretation and critical analysis, and focusing on areas that require human expertise, such as patient counseling and ethical decision-making. Ultimately, the successful integration of LLMs in pharmacy practice holds great potential to enhance patient care, foster innovation, and advance the field of pharmaceutical sciences.

## Notes

### Competing Interest Statement

The authors have declared no competing interest.

